# Functional dissection of KATP channel structures reveals the importance of a conserved interface

**DOI:** 10.1101/2023.08.03.551891

**Authors:** Yaxiong Yang, Lei Chen

**Author notes:** Corresponding author: Lei Chen.

## Abstract

ATP-sensitive potassium channels (KATP) are inhibited by ATP but activated by Mg-ADP, coupling the intracellular ATP/ADP ratio to the potassium conductance of the plasma membrane. Although there has been progress in determining the structure of KATP channels, the functional significance of the domain-domain interface in the gating properties of KATP channels is not fully understood. In this study, we propose a new two-module assembly model for the KATP channel. Our mutagenesis experiments, based on this model, indicate that deleting ECL3 on the SUR1 subunit impairs KNtp-independent Mg-ADP activation. This finding demonstrates the essential role of intramolecular interactions between KATP_core_ and SUR_ABC_ in Mg-ADP activation. Notably, this interface is functionally conserved between SUR1 and SUR2. Additionally, the hydrophobic residue F351 on ECL3 of SUR1 is crucial for maintaining the stability of this interface.

## Introduction

KATP channels are metabolic sensors on the plasma membrane. Their potassium currents are inhibited by intracellular ATP and activated by intracellular Mg-ADP. Therefore, they tune the membrane potential of the cell according to the intracellular ATP/ADP ratio (Nichols, 2006). KATP channels are distributed in many tissues, such as pancreatic cells, the brain, and muscles, and are involved in a series of physiological processes (Ashcroft, 2006; Flagg et al., 2010). The genetic mutations of KATP channels can lead to various human diseases, including dilated cardiomyopathy (Bienengraeber et al., 2004), Cantu syndrome (Harakalova et al., 2012; van Bon et al., 2012), familial atrial fibrillation (Olson et al., 2007), intellectual disability myopathy syndrome (Smeland et al., 2019), neonatal diabetes mellitus (Pipatpolkai et al., 2020), and hyperinsulinaemic hypoglycemia of infancy (Pipatpolkai et al., 2020). KATP channel inhibitors (insulin secretagogues) are widely used to treat diabetes. KATP activators (KATP openers) are used for the treatment of hypoglycemia, hypertension, and hair loss (Wu et al., 2022).

KATP channels are hetero-octamers formed by Kir6 (Kir6.1 or Kir6.2) and SUR (SUR1, SUR2A, or SUR2B) subunits in a 4:4 ratio. Four Kir6 subunits form the central channel pore, and four regulatory SUR subunits are located peripherally (Wu et al., 2022). Recent cryo-EM studies have provided high-resolution views of the pancreatic KATP channel (Kir6.2+SUR1) in multiple states (Ding et al., 2019; Lee et al., 2017; Li et al., 2017; Martin et al., 2017a; Martin et al., 2019; Martin et al., 2017b; Wang et al., 2022a; Wang et al., 2022b; Wu et al., 2018; Zhao and MacKinnon, 2021). These structures show that in the inhibited condition, when insulin secretagogues and ATP are present, the ATP binds in a pocket of Kir6.2, and the ABLOS motif on SUR1 interacts with the phosphate group of ATP (Ding et al., 2019). The Kir6 N-terminal peptide (KNtp) binds in the center of SUR1, which is close to the insulin secretagogue binding site (Ding et al., 2019; Martin et al., 2019; Wu et al., 2018). In the activated condition, when Mg-ADP is present, Mg-ADP binds in the consensus site of NBD2 of SUR1 to drive the asymmetric dimerization of the ABC-transporter module (Lee et al., 2017; Wang et al., 2022b; Wu et al., 2018; Zhao and MacKinnon, 2021) and to release the KNtp from SUR1 (Wang et al., 2022b; Wu et al., 2018; Zhao and MacKinnon, 2021). The unbinding of ATP from Kir6.2 is associated with the rotation of Kir6.2 CTD, leading to channel opening (Wang et al., 2022b; Zhao and MacKinnon, 2021). Despite the progress made, there is still incomplete understanding of how the ligand binding on SUR regulates the activity of the Kir6 channel. Structural studies have predominantly revealed a propeller architecture of the pancreatic KATP channel. However, it is worth noting that one study observed both the propeller form and the quatrefoil form of the channel coexisting within a single sample. (Lee et al., 2017). Strikingly, the same phenomenon was observed in the vascular KATP channel in the inhibitory condition (Kir6.1+SUR2B) (Sung et al., 2021). The transition from the propeller form to the quatrefoil form of the SUR subunits involves significant structural changes and rearrangements of the inter-domain interfaces. However, the biological significance of these two distinct architectures, particularly the quatrefoil form, is still largely unknown. Further research is needed to uncover the functional implications and role of these different structural conformations in the activity of the KATP channel.

In our study, we propose a revised two-module model for the assembly of KATP channels. To investigate the role of inter-module interfaces in the inhibition and activation of KATP channels, we employ structure-based site-directed mutagenesis along with electrophysiology experiments. By manipulating and studying these specific interfaces, we aim to gain a better understanding of how they contribute to the regulation and function of KATP channels.

## Results

### Two-module model for KATP channel

The available structural data for both the propeller and the quatrefoil forms of KATP channels indicate that the Kir6 subunit and TMD0 domains of SUR are tightly associated through a stable interface (Lee et al., 2017; Sung et al., 2021). However, there are significant differences in the relative orientations between the TMD0 domain of SUR and the remaining ABC transporter module between the propeller and the quatrefoil structures (Lee et al., 2017; Sung et al., 2021). Based on these observations, we propose that the Kir6 and TMD0 domains of SUR (1-214 of SUR1 or 1-212 of SUR2B) form a stable core module known as KATP_core_. This core module has a dynamic interface with the remaining ABC transporter module (SUR_ABC_) of SUR (TMD1-NBD1-TMD2-NBD2, 215-C terminus of SUR1 or 213-C terminus of SUR2B) (Fig. 1a-c).

**Fig. 1.**
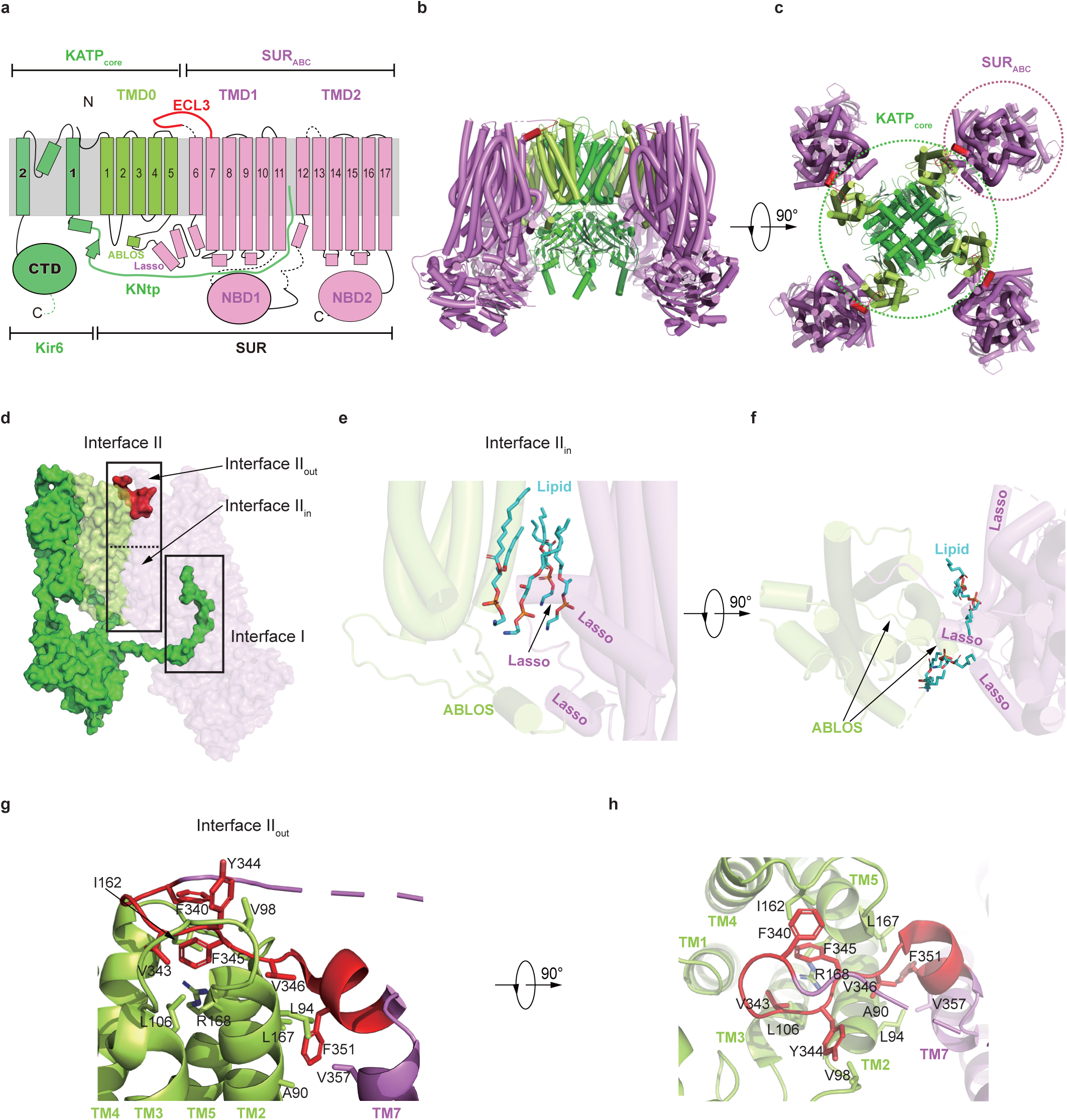
The two interfaces between KATP_core_ and SUR_ABC_. **A**Topology of a KATP channel consisting of Kir6 and SUR. TMD, transmembrane domain; ECL, extracellular loop; NBD, nucleotide-binding domain; CTD, cytoplasmic domain; KNtp, Kir6 N-terminal peptide; ABLOS, ATP-binding loop on SUR. The Kir6 and TMD0 of SUR form the KATP_core_, and the TMD1-NBD1-TMD2-NBD2 of SUR form the SUR_ABC_. Kir6, TMD0, and TMD1-NBD1-TMD2-NBD2 are colored in green, lemon, and violet, respectively. The ECL3 and KNtp are in bold and colored in red and green, respectively. All transmembrane helices are shown as cylinders, and the phospholipid bilayer is represented using gray area. **b** The side view of a tetrameric KATP structure (PDB ID: 7W4P). **c** The top view of the tetrameric KATP structure (PDB ID: 7WAP). The KATP_core_ and SUR_ABC_ are circled in dashes. **d** The side view of a KATP structure (one Kir6.1 subunit and one SUR2B subunit, PDB ID: 7MIT). The ECL3 and Kir6 are shown as non-transparent surfaces, colored in red and green, respectively. The SUR is shown as a semitransparent surface. The Interface I and Interface II between KATP_core_ and SUR_ABC_ are boxed. The intracellular region and extracellular region of the Interface II are denoted as Interface II_in_ and Interface II_out_, respectively. **e** Close-up view of Interface II_in_ in the KATP structure (PDB ID: 6JB1). The lipid molecules are shown as sticks and colored in blue. The lasso motif and ABLOS motif are denoted. **f** The top view of Interface II_in_. **g** Overview of the interactions in the Interface II_out_ of pancreatic KATP (PDB ID: 7W4P). The side chains of residues that are involved in the interactions between the ECL3 of the SUR_ABC_ and the groove of TMD0 of KATP_core_ are shown as sticks. **h** The top view of the interactions in Interface II_in_.

Based on our newly proposed assembly model, we have identified two interfaces between KATP_core_ and SUR_ABC_ in the propeller structure (Fig. 1d). Interface I involves the interactions between KNtp in KATP_core_ and SUR_ABC_ (Fig. 1d). This interface was initially discovered in the pancreatic KATP channel (Wu et al., 2018) and later observed in both the propeller and quatrefoil forms of the vascular KATP channel (Sung et al., 2021). It is important to note that the existence of Interface I is state-dependent. KNtp binds within the cytoplasmic central vestibule of SUR_ABC_, and the binding site is accessible only when SUR_ABC_ is in the inward-facing conformation (Martin et al., 2019; Sung et al., 2021; Wu et al., 2018). However, this binding site is disrupted when SUR_ABC_ transitions to the occluded conformation in the presence of Mg-ADP, which occurs in both SUR1 (Lee et al., 2017; Sung et al., 2021; Wang et al., 2022b; Wu et al., 2018) and SUR2 (Ding et al., 2022). Indeed, the state-dependent interaction between KNtp and SUR_ABC_ correlates with the significant impact observed upon the deletion of KNtp on ATP-sensitivity (Babenko et al., 1999; Koster et al., 1999; Reimann et al., 1999), as well as the binding and inhibition of insulin secretagogues (Devaraneni et al., 2015; Kuhner et al., 2012; Reimann et al., 1999). These studies have provided evidence for the functional importance of KNtp in modulating the ATP-sensitivity and response to insulin secretagogues of KATP channels.

Interface II is situated between SUR_ABC_ and TMD0 of SUR1 in the KATP_core_ (Fig. 1d). This interface can be divided into two portions: Interface II_in_, which is located in the intracellular region, and Interface II_out_, which is situated in the extracellular region. In the structure of pancreatic KATP in complex with repaglinide (PDB ID: 6JB1) (Ding et al., 2019), Interface II_in_ is well resolved and shows the involvement of multiple lipid molecules between KATP_core_ and SUR_ABC_ (Fig. 1e, f). This suggests that the stability of Interface II_in_ may be influenced by the lipidic environment and could be sensitive to harsh detergent treatments. On the other hand, Interface II_out_ is best resolved in the structure of KATP with the H175K mutant on Kir6.2 (PDB ID: 7W4P) (Wang et al., 2022b). Within Interface II_out_, the extracellular loop 3 (ECL3) between TM6 and TM7 of SUR_ABC_ binds within a groove of TMD0 in the KATP_core_. Specifically, V346 and F351 on ECL3, along with V357 on TM7, establish hydrophobic interactions with A90 and L94 on TM2, as well as L167 on TM5 of TMD0. F345 on ECL3 forms a cation-π interaction with R168 on TM5. Moreover, F340, V343, and Y344 contribute to hydrophobic interactions with I162 on ECL2, L106 on TM3, and V98 on ECL1, respectively (Fig. 1g, h). In the quatrefoil structure, Interface II is completely disrupted. This indicates a significant conformational change and loss of the interactions that normally occur at Interface II in propeller form. The disruption of Interface II in the quatrefoil form of the KATP channel suggests that this interface is not present or functional in this particular conformation.

### SUR1 mutant with ECL3 deletion could switch between states in a ligand-dependent manner

To investigate the function of Interface II, we conducted a deletion of ECL3 in SUR1 (residues 340-352 of SUR1) to create SUR1_ΔECL3_. This deletion was designed to destabilize the extracellular portion of Interface II. We then examined whether SUR1_ΔECL3_ was properly folded and capable of transitioning between the inward-facing conformation and the occluded conformation, similar to SUR1_WT_. Since glibenclamide binds to SUR1 in the inward-facing conformation, leading to the recruitment of KNtp and inhibition of KATP currents in the absence of ATP (Wu et al., 2018), we utilized glibenclamide inhibition to assess whether SUR1_ΔECL3_ was properly folded and capable of adopting the inward-facing conformation.

The co-expression of Kir6.2 with SUR1_ΔECL3_ resulted in the generation of robust ATP-sensitive potassium currents (Fig. 2a, b). This indicates that SUR1_ΔECL3_ is capable of assembling with Kir6.2 and facilitating the trafficking of Kir6.2 to the plasma membrane. Furthermore, inside-out recordings demonstrated that glibenclamide was able to inhibit the current mediated by the KATP channel formed by SUR1_ΔECL3_ (Fig. 2c, d). This suggests that SUR1_ΔECL3_ is properly folded and retains both the glibenclamide-binding site and the KNtp-binding site, both of which are crucial for glibenclamide inhibition.

**Fig. 2.**
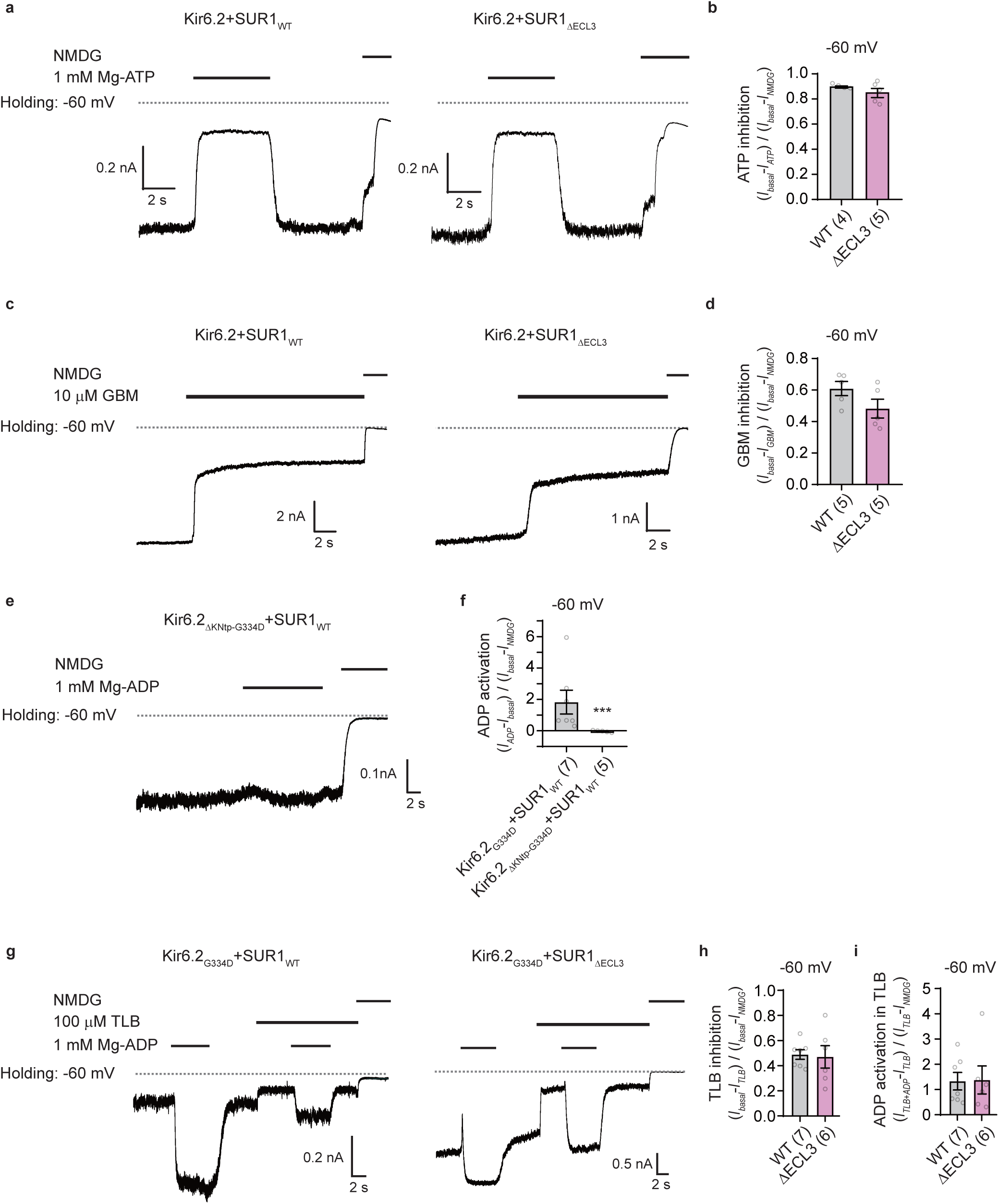
Functional analyses of the Interface II in the SUR by electrophysiology. **a**Representative inside-out electrophysiological recordings of K_ATP_ channels formed by Kir6.2 and SUR1 wild type (SUR1_WT_), or truncated SUR1 which lacks ECL3 (SUR1_ΔECL3_). 1 mM Mg-ATP is applied to inhibit the channels. And the bath solution containing the impermeant cation NMDG (N-methyl-d-glucamine) is applied to determine the leakage of the patch clamp. The holding potential is −60 mV. **b** Statistical summary of the 1 mM Mg-ATP inhibition. ATP inhibition is calculated by the ratio between the ATP-inhibited currents (*I_basal_* - *I_ATP_*) and the corrected basal currents (*I_basal_* – *I_NMDG_*). **c** Representative traces of K_ATP_ channels formed by Kir6.2 and SUR1_WT_, or SUR1_ΔECL3_. 10 μM insulin secretagogues, GBM (glibenclamide), is applied to inhibit the channel. And NMDG is applied to determine the leak currents. The holding potential is −60 mV. **d** A statistical summary of the 10 μM GBM inhibition. GBM inhibition is calculated by the ratio between GBM-inhibited currents (*I_basal_*– *I_GBM_*) and the corrected basal currents (*I_basal_*– *I_NMDG_*). **e** Representative traces of the KATP channel formed by Kir6.2_ΔKNtp-G334D_ and SUR1_WT_. The holding potential is −60 mV. 1 mM Mg-ADP is applied to activate the channel and NMDG is applied to determine the leakage. **f** Statistical comparison of the 1 mM Mg-ADP activation between the KATP channel formed by Kir6.2_G334D_ and SUR1_WT_, and the KATP channel formed by Kir6.2_ΔKNtp-G334D_ and SUR1_WT_. The ADP activation is calculated by the ratio between the ADP-activated currents (*I_ADP_* - *I_basal_*) and the corrected basal currents (*I_basal_*– *I_NMDG_*). **g** Representative traces of KATP channels formed by Kir6.2_G334D_ and SUR1_WT_, or SUR1_ΔECL3_. The holding potential is −60 mV. The insulin secretagogues TLB (tolbutamide) and Mg-ADP are applied to inhibit and activate the channel, respectively. NMDG is applied to determine the leakage. **h** Statistical summary of TLB inhibition. The ratio between downregulated currents by TLB (*I_basal_*– *I_TLB_*) and corrected basal currents (*I_basal_*– *I_NMDG_*) serves as the index. **i** Statistical summary of Mg-ADP activation under TLB treatment. The ratio between upregulated currents by Mg-ADP under TLB (*I_TLB+ADP_*– *I_TLB_*) and the currents in TLB (*I_TLB_* – *I_NMDG_*) serves as the index for ADP activation. All data are shown as mean ± SEM. Two-tailed unpaired Student’s *t*-test was calculated for (**f**) with criteria of significance; *** *P*□<□0.001.

To investigate whether SUR1_ΔECL3_ could adopt the occluded conformation, we utilized the G334D mutant of Kir6.2 (Kir6.2_G334D_), which abolishes the inhibitory binding of ATP and ADP on Kir6.2 (Proks et al., 2010) to only show the activation effect of Mg-ADP. In experiments conducted with SUR1_WT_ and Kir6.2_G334D_, the KATP channel was found to be activated by Mg-ADP, primarily due to the release of KNtp from its binding site. This activation by Mg-ADP was largely abolished upon further deletion of KNtp (Fig. 2e, f). Notably, this Mg-ADP activation was sensitive to the low-affinity insulin secretagogue, tolbutamide (Fig. 2g-i). Similarly, in our investigation, we observed that the KATP channel formed by Kir6.2_G334D_ and SUR1_ΔECL3_ could also be activated by Mg-ADP and subsequently inhibited by tolbutamide (Fig. 2g-i). These findings provide evidence that SUR1_ΔECL3_ is capable of adopting the occluded conformation in the presence of Mg-ADP, similar to SUR1_WT_. These collective findings support the notion that SUR1_ΔECL3_ is well-folded and capable of dynamic conformational changes, transitioning between the inward-facing and occluded conformations, in response to ligand binding.

### Deletion of ECL3 in SUR1 abolishes Mg-ADP activation in the absence of KNtp

The importance of Interface I for the coupling between KATP_core_ and SUR_ABC_ has been established. To further investigate the function of Interface II, we deliberately removed KNtp from Kir6.2 to disrupt Interface I. In inside-out recordings, it was observed that in the absence of KNtp, Mg-ADP robustly reactivated ATP-inhibited KATP currents (Fig. 3a, b). This suggests that Interface II is involved in KNtp-independent Mg-ADP activation. These findings align with previous recordings conducted on the SUR1-Kir6.2 fusion protein, where the covalent linkage between the C-terminus of SUR1 and KNtp essentially abolished Interface I (Ding et al., 2019; Wu et al., 2018). Additionally, the Mg-ADP activation can be slightly potentiated by the SUR1 activator NN414 (Fig. 3c, d). These results provide further support for the functional role of Interface II in mediating KNtp-independent Mg-ADP activation.

**Fig. 3.**
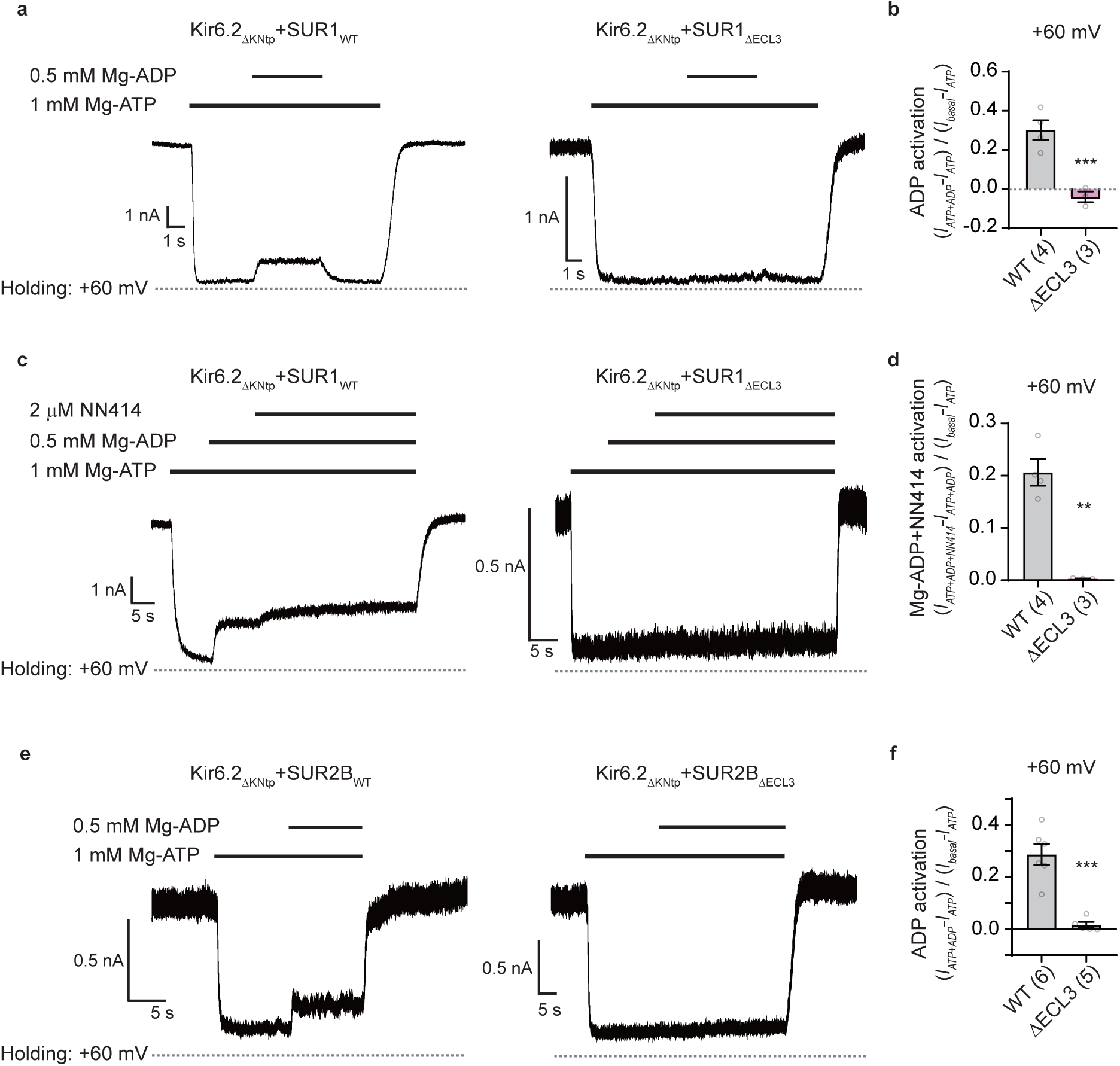
The role of ECL3 in Mg-ADP activation of KATP channels. **a** Representative inside-out electrophysiological recordings of K_ATP_ channels formed by Kir6.2_ΔKNtp_ and SUR1_WT_, or SUR1_ΔECL3_. The holding potential is +60 mV. 1 mM Mg-ATP and an additional 0.5 mM Mg-ADP are applied for inhibition and re-activation of the channels, respectively. **b** Statistical summary of ADP activation for SUR1_WT_ and SUR1_ΔECL3_. The ratio between upregulated currents by Mg-ADP (*I_ATP+ADP_* - *I_ATP_*) and basal currents (*I_basal_* - *I_ATP_*) serves as the index. **c** Representative traces of K_ATP_ channels formed by Kir6.2_ΔKNtp_ and SUR1_WT_ or SUR1_ΔECL3_. The holding potential is +60 mV. 2 μM NN414 is applied to the bath solution containing 1 mM Mg-ATP and 0.5 mM Mg-ADP to test the activation effect. **d** Statistical summary of NN414 activation upon 1 mM Mg-ADP and 0.5 mM Mg-ADP for SUR1_WT_ and SUR1_ΔECL3_. The ratio between upregulated currents by NN414 (*I_ATP+ADP+NN414_*-*I_ATP+ADP_*) and basal currents (*I_basal_* - *I_ATP_*) serves as the index. **e** Representative traces of KATP channels formed by Kir6.2_ΔKNtp_ and SUR2_WT_, or SUR2B_ΔECL3_ for testing of 0.5 mM Mg-ADP activation under 1 mM Mg-ATP inhibition. The holding potential is +60 mV. **f** Statistical summary of ADP activation for the SUR2B_WT_ and SUR2B_ΔECL3_. All data are shown as mean ± SEM. A two-tailed unpaired Student’s *t*-test was calculated for (**b**), (**d**) and (**f**) with criteria of significance; ** *P* < 0.01; *** *P*□<□0.001.

Interestingly, when ECL3 on Interface II was deleted, we observed the complete abolition of re-activation by both Mg-ADP alone and Mg-ADP in combination with NN414 (Fig. 3a-d). These results strongly indicate that Interface II plays a crucial role in transmitting the activation signal from Mg-ADP binding on SUR_ABC_ to KATP_core_. Furthermore, we discovered a similar phenomenon when ECL3 was deleted in SUR2B (Fig. 3e, f), suggesting that the function of Interface II is conserved between both SUR1 and SUR2. These findings highlight the importance of Interface II in mediating the transmission of the Mg-ADP activation signal and further support its role in KATP channel regulation across different SUR isoforms.

### F351 on SUR1 is important for the stability of Interface II

Sequence alignments clearly demonstrate that the length of ECL3 varies between SUR1 and SUR2, particularly in the region between residues 340 and 346 of SUR1 (Fig. 4a), suggesting this region is not essential for binding to TMD0. Indeed, we found that when coexpressing with Kir6.2_ΔKNtp_, SUR1_Δ340-346_ exhibited normal Mg-ADP activation (Fig. 4b). In stark contrast, the Mg-ADP activation of SUR1Δ347-352 was impaired, similar to the phenotype observed for SUR1_ΔECL3_ (Fig. 4c). This suggests that residues within the 347-352 region of ECL3 are crucial for binding with TMD0. Additionally, through sequence alignment, we observed that several residues within the 347-352 region are conserved between SUR1 and SUR2 (Fig. 4a). Among these residues, the side chain of F351 is involved in hydrophobic interactions, indicating its potential significance in the interaction between ECL3 and TMD0. To disrupt the side chain-involving interactions within Interface II, we introduced a mutation in which F351 was changed to glycine, resulting in the generation of SUR1_F351G_. Upon co-expression of SUR1_F351G_ with Kir6.2_ΔKNtp_, it was observed that the Mg-ADP reactivation of ATP-inhibited currents was largely abolished, closely resembling the effect observed with ECL3 deletion (Fig. 4d). This suggests that F351 is crucial for maintaining the stability and function of Interface II (Fig. 4e). Remarkably, when SUR1_F351G_ was co-expressed with Kir6.2_G334D_, it conferred Mg-ADP activation that could be inhibited by tolbutamide (Fig. 4f-h). This indicates that SUR1_F351G_ is folded properly and capable of transitioning between the inward-facing and occluded states in response to ligand binding. Collectively, these pieces of evidence strongly support the significance of the conserved F351 residue within Interface II and highlight its vital role in the proper functioning of this interface.

**Fig. 4.**
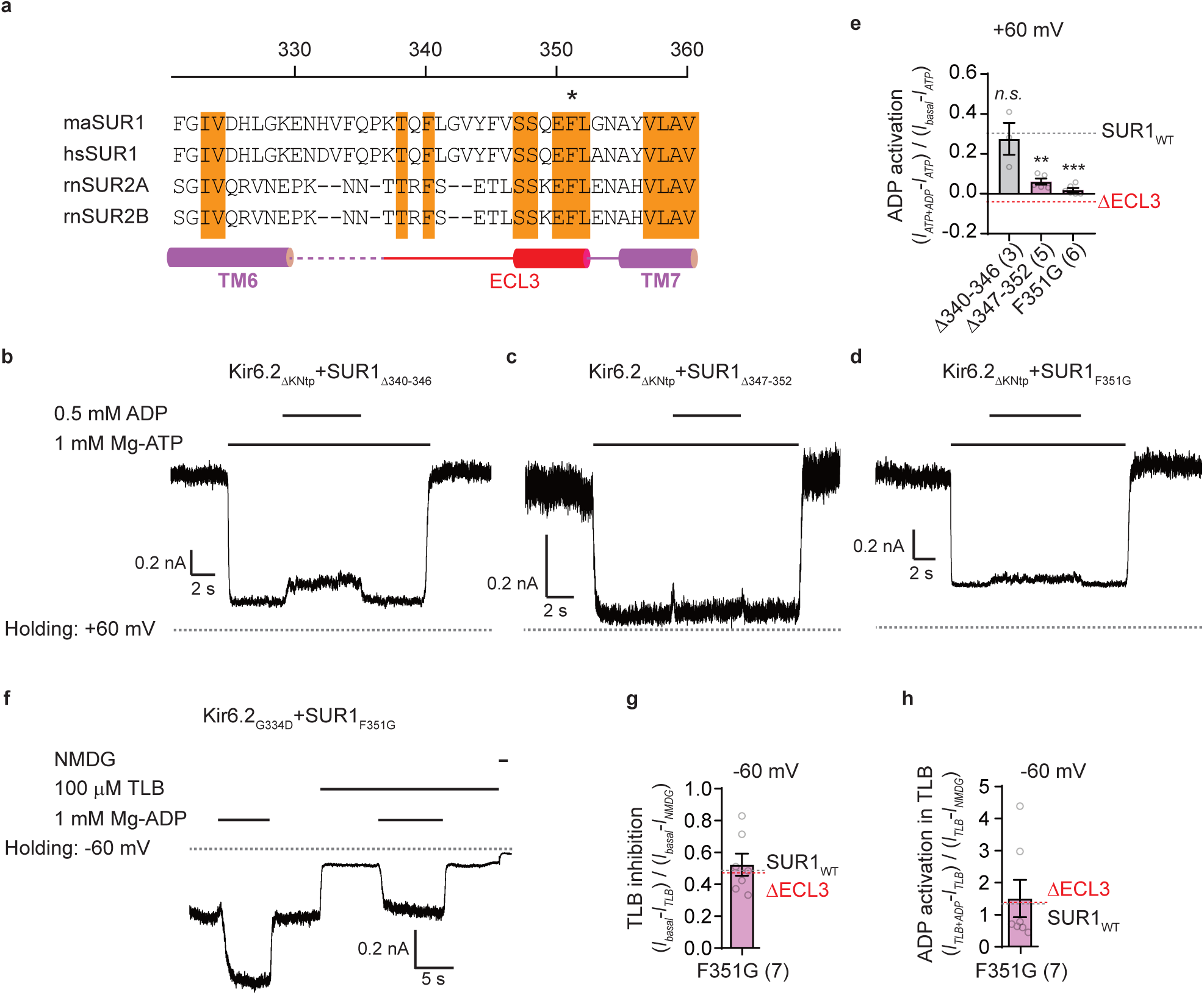
F351 of SUR1 is important for the interactions between ECL3 and TMD0. **a** Sequence alignment of TM6-ECL3-TM7 across *Mesocricetus auratus* SUR1 (maSUR1), *Homo sapiens* SUR1 (hsSUR1), *Rattus norvegicus* SUR2A (rnSUR2A), and *Rattus norvegicus* SUR2B (rnSUR2B). Secondary structure elements for TM6-ECL3-TM7 in maSUR1 are shown as cylinders (helices) and lines (loops). The TM6 and TM7 are colored in violet, and the ECL3 is colored in red. Unmodeled residues are shown as dashed lines. Identical amino acids are highlighted in orange. The key residue of F351 is marked with an asterisk above. **b-d** The SUR1_Δ340-346_, SUR1_Δ347-352_ and SUR1_F351G_ are co-expressed with Kir6.2_ΔKNtp_ for testing of 0.5 mM Mg-ADP activation under 1 mM Mg-ATP. The holding potential is +60 mV. **e** Statistical summary of Mg-ADP activation under Mg-ATP treatment. Gray and pink dotted lines representing the mean value of SUR1_WT_ and SUR1_ΔECL3_ from Fig. 3b are denoted for comparison. **f** Representative traces of K_ATP_ channels formed by Kir6.2_ΔKNtp_ and SUR1_F351G_ for testing of 100 μM TLB inhibition and 1 mM Mg-ADP activation. NMDG is applied to determine the leakage. The holding potential is −60 mV. **g** Statistical summary of TLB inhibition. Gray and pink dotted lines representing the mean values of SUR1_WT_ and SUR1_ΔECL3_ from Fig. 2h are shown for comparison with SUR1_F351G_. **h** Statistical summary of Mg-ADP activation under TLB treatment. The gray and pink dotted lines are from Fig. 2i for comparison. All data are shown as mean ± SEM. One-way ANOVA followed by Dunnett post hoc tests were used for (**e**) with the control group of SUR1_WT_. Criteria of significance; ** *P* < 0.01; *** *P*□<□0.001.

## Discussions

### The propeller structure versus the quatrefoil structure of KATP

In this study, we aimed to destabilize Interface II by deleting ECL3 in SUR. Although the extent to which the deletion of ECL3 disrupts Interface II remains unknown, the functional impairment observed in the ECL3-deletion mutant clearly highlights the significance of Interface II in KNtp-independent Mg-ADP activation. The complete disruption of Interface II in the quatrefoil structure suggests that KATP in this form is defective in KNtp-independent Mg-ADP activation. Interestingly, our results demonstrate that the ECL3-deletion mutants retain KNtp-dependent glibenclamide inhibition, indicating that Interface II is not essential for this process. This finding is consistent with the fact that KNtp binds to the central vestibule of vascular KATP in both the propeller and quatrefoil forms, indicating the integrity of Interface I in both forms. It should be noted that the currently available KATP structures are derived from detergent-solubilized KATP proteins, and further investigation is required to determine the proportion of the quatrefoil KATP that exists in vivo on the cell membrane.

### The function of Interface I and Interface II of KATP

KATP channels are fascinating allosteric protein complexes that arise from the association of the Kir6 ion channel with the SUR ABC transporter. Previous research has demonstrated the existence of functional coupling between Kir6 and SUR, whereby ligand binding on SUR impacts the activity of Kir6. This coupling is evident in the unique gating characteristics exhibited by KATP channels, including ATP inhibition, insulin secretagogue inhibition, and Mg-ADP activation. Understanding the communication between SUR and Kir6 is a fundamental question in the field. In this study, we redefine the structural organization of the KATP channel as a two-module complex composed of KATP_core_ and SUR_ABC_. Within this complex, two physical interactions, referred to as Interface I and Interface II, are observed in propeller structures. In the following discussion, we aim to associate the known functional properties of KATP with the structural changes occurring at Interface I and Interface II.

The ATP inhibition property is inherent to the Kir6 subunit itself (Tucker et al., 1997). However, when the SUR subunit is co-expressed, it enhances the ATP sensitivity of Kir6 (Aittoniemi et al., 2009). This enhancement is partially attributed to the direct interaction between ATP and the K205 residue on the ABLOS motif of SUR (Ding et al., 2019). The ABLOS motif is located near Interface II and undergoes a conformational change upon binding of Mg-ADP to SUR_ABC_ (Wang et al., 2022b). Moreover, the KNtp region of the KATP_core_ binds to the central vestibule of SUR_ABC_ (Interface I), resulting in increased ATP affinity (Babenko et al., 1999; Koster et al., 1999; Reimann et al., 1999). This binding likely allosterically stabilizes Kir6 in a closed conformation that exhibits a high affinity for ATP. Furthermore, structural studies have observed the binding of KNtp to SUR_ABC_ in the presence of ATP (Martin et al., 2019), providing further support for this mechanism.

The activation of KATP by Mg-ADP involves two distinct processes: a KNtp-dependent process and a KNtp-independent process. The KNtp-dependent activation can be clearly observed in the Mg-ADP activation of the channel formed by Kir6.2_G334D_ (Fig. 2g). However, when KNtp is further deleted, this activation is completely abolished (Fig. 2e). Structurally, the dimerization of SUR_ABC_ induced by Mg-ADP disrupts Interface I and causes the expulsion of KNtp from its binding site (Wu et al., 2018). The Mg-ADP activation of Kir6.2_G334D_ suggests that, in the absence of ATP, a fraction of the KATP channel has KNtp bound to SUR_ABC_. However, the binding of KNtp in SUR1_ABC_ was not observed in the structure of KATP in the apo state (Martin et al., 2019), possibly due to the low occupancy of KNtp. To detect the presence of the small fraction of SUR1 with KNtp bound in the apo state, a more thorough 3D classification approach might be required during the image processing stage. When Interface I is completely disrupted in the KNtp deletion mutant, Mg-ADP can reactivate the ATP-inhibited currents of KATP (Fig. 3a), indicating the involvement of Interface II in this process. From a structural perspective, the dimerization of SUR1_ABC_ induced by Mg-ADP binding leads to an outward movement of the ABLOS motif on KATP_core_. This conformational change might result in a reduced affinity for ATP and subsequent reactivation of the channel (Wang et al., 2022b). In the presence of both the KNtp deletion and the G334D mutation, Mg-ADP activation was not observed anymore (Fig. 2e). This suggests that KNtp-independent Mg-ADP activation operates by reducing the affinity of ATP, rather than directly enhancing the activity of KATP_core_.

KATP activity can be inhibited by insulin secretagogues, which bind to SUR_ABC_. This inhibition can be divided into two distinct processes: direct inhibition in the absence of nucleotides and inhibition of Mg-ADP activation. It has been demonstrated that the direct inhibition is mediated by Interface I, as both the deletion of KNtp (Devaraneni et al., 2015; Kuhner et al., 2012; Reimann et al., 1999) and the fusion of KNtp to the C-terminus of SUR (Ding et al., 2019; Wu et al., 2018) abolish this type of inhibition. On the other hand, the inhibition of Mg-ADP activation in the absence of KNtp (Fig. 2g) primarily occurs because the binding of insulin secretagogues to SUR_ABC_ prevents the transition of SUR_ABC_ to the occluded state, thus blocking the signaling through Interface II.

In summary, the underlying structural mechanisms of three hallmarks of KATP channels, namely the inhibition by ATP, the activation by Mg-ADP, and the inhibition by insulin secretagogues, all mechanistically relate to not only Interface I but also Interface II. The concerted structural changes in these two interfaces shape the overall gating properties of the KATP channel.

## Methods

### Cell lines

FreeStyle 293F (Thermo Fisher Scientific # R79007) suspension cells were cultured in 293 Expression Medium (Gibco) supplemented with 1% FBS at 37□°C in CO_2_ incubator. Cell lines were free of mycoplasma contamination, tested by MycoBlue Mycoplasma Detector assay (Vazyme, D101-02).

### Constructs

cDNA of Kir6.2 was from *Mus musculus* and cloned into a customized pBM BacMam vector with a C-terminal GFP tag. cDNA of SUR1 and SUR2B were from *Mesocricetus auratus* and *Rattus norvegicus*, respectively and were cloned into a customized pBM BacMam expression vector without the GFP tag(Ding et al., 2023; Wang et al., 2022b). All deletions and point mutations were followed by the site-directed mutagenesis with PrimeSTAR DNA Polymerase (Takara Bio).

### Electrophysiology

KATP constructs (Kir6.2-CGFP and SUR) were co-transfected into FreeStyle 293-F cells using polyethylenimine at a cell density of 1□×□10^6^ cells/ml. Cells were cultured in FreeStyle 293 Expression Medium with 1% FBS for 1 day before electrophysiological recording. Macroscopic currents were recorded using inside-out mode at the holding potential of +60□mV or −60 mV in the pipette (membrane potential of −60□mV or +60 mV) through an Axon-patch 200B amplifier (Axon Instruments, USA). Patch electrodes were pulled by a horizontal micro-electrode puller (P-1000, Sutter Instrument Co, USA) and heat-polished by a pipette microforge to adjust the resistances to around 1-3 MΩ. Both pipette and bath solutions were KINT buffer, containing (mM): 140 KCl, 1 EGTA, 2 MgCl_2_, and 10 HEPES (pH 7.4, adjusted by KOH). The NMDG, bath solution containing (mM): 140 NMDG, 1 EGTA, and 10 HEPES, was applied to determine the leakage of patch. Signals were acquired at 5□kHz and low-pass filtered at 1□kHz. Data were further analyzed with Clampfit, Excel, and GraphPad.

### Figure preparation

KATP structures are from PDB ID: 7W4P, 7MIT and 6JB1. All images of KATP structures were prepared using the PyMOL 2.4.1.

### Statistics and reproducibility

Data were analyzed in GraphPad Prism software, and shown as mean□±□SEM (Standard Error of the Mean). Two-tailed unpaired Student’s *t*-test was performed to compare two groups. One-way ANOVA followed by Dunnett for post hoc tests were performed to compare more than two groups. Criteria of significance; ** *P* < 0.01; *** *P*□<□0.001. No less than 3 independent patches are recorded for each condition.

## Acknowledgement

This work is the result of extensive collaboration and fruitful discussions among members of the KATP subgroup in Chen Lab, including Jing-Xiang Wu, Dian Ding, Mengmeng Wang, and Tianyi Hou. The work is supported by grants from the Ministry of Science and Technology of China (National Key R&D Program of China, 2022YFA0806504 to L.C.), the National Natural Science Foundation of China (91957201, 32225027, and 31821091 to L.C., and 32271015 to Y.Y.) the Fundamental Research Funds for the Central Universities (YWF-21-BJ-J-1150 to Y.Y.), and Center For Life Sciences (CLS to L.C.).

## Contributions

L.C. initiated the project and wrote the manuscript draft. Y.Y. carried out the experiments and prepared figures. Both authors contributed to the manuscript preparation.

## Reference

Aittoniemi, J., Fotinou, C., Craig, T.J., de Wet, H., Proks, P., and Ashcroft, F.M. (2009). Review. SUR1: a unique ATP-binding cassette protein that functions as an ion channel regulator. Philos. Trans. R. Soc. Lond. B. Biol. Sci. 364, 257–267.

Ashcroft, F.M. (2006). K(ATP) channels and insulin secretion: a key role in health and disease. Biochem. Soc. Trans. 34, 243–246.

Babenko, A.P., Gonzalez, G., and Bryan, J. (1999). The N-terminus of KIR6.2 limits spontaneous bursting and modulates the ATP-inhibition of KATP channels. Biochem. Biophys. Res. Commun. 255, 231–238.

Bienengraeber, M., Olson, T.M., Selivanov, V.A., Kathmann, E.C., O’Cochlain, F., Gao, F., Karger, A.B., Ballew, J.D., Hodgson, D.M., Zingman, L.V., et al. (2004). ABCC9 mutations identified in human dilated cardiomyopathy disrupt catalytic KATP channel gating. Nat. Genet. 36, 382–387.

Devaraneni, P.K., Martin, G.M., Olson, E.M., Zhou, Q., and Shyng, S.L. (2015). Structurally distinct ligands rescue biogenesis defects of the KATP channel complex via a converging mechanism. J. Biol. Chem. 290, 7980–7991.

Ding, D., Hou, T., Wei, M., Wu, J.X., and Chen, L. (2023). The inhibition mechanism of the SUR2A-containing K(ATP) channel by a regulatory helix. Nat Commun 14, 3608.

Ding, D., Wang, M., Wu, J.X., Kang, Y., and Chen, L. (2019). The Structural Basis for the Binding of Repaglinide to the Pancreatic KATP Channel. Cell Rep 27, 1848–1857 e1844.

Ding, D., Wu, J.X., Duan, X., Ma, S., Lai, L., and Chen, L. (2022). Structural identification of vasodilator binding sites on the SUR2 subunit. Nat Commun 13, 2675.

Flagg, T.P., Enkvetchakul, D., Koster, J.C., and Nichols, C.G. (2010). Muscle KATP channels: recent insights to energy sensing and myoprotection. Physiol. Rev. 90, 799–829.

Harakalova, M., van Harssel, J.J., Terhal, P.A., van Lieshout, S., Duran, K., Renkens, I., Amor, D.J., Wilson, L.C., Kirk, E.P., Turner, C.L., et al. (2012). Dominant missense mutations in ABCC9 cause Cantu syndrome. Nat. Genet. 44, 793–796.

Koster, J.C., Sha, Q., and Nichols, C.G. (1999). Sulfonylurea and K(+)-channel opener sensitivity of K(ATP) channels. Functional coupling of Kir6.2 and SUR1 subunits. J. Gen. Physiol. 114, 203–213.

Kuhner, P., Prager, R., Stephan, D., Russ, U., Winkler, M., Ortiz, D., Bryan, J., and Quast, U. (2012). Importance of the Kir6.2 N-terminus for the interaction of glibenclamide and repaglinide with the pancreatic K(ATP) channel. Naunyn. Schmiedebergs Arch. Pharmacol. 385, 299–311.

Lee, K.P.K., Chen, J., and MacKinnon, R. (2017). Molecular structure of human KATP in complex with ATP and ADP. Elife 6.

Li, N., Wu, J.X., Ding, D., Cheng, J., Gao, N., and Chen, L. (2017). Structure of a Pancreatic ATP-Sensitive Potassium Channel. Cell 168, 101–110 e110.

Martin, G.M., Kandasamy, B., DiMaio, F., Yoshioka, C., and Shyng, S.L. (2017a). Anti-diabetic drug binding site in a mammalian KATP channel revealed by Cryo-EM. Elife 6.

Martin, G.M., Sung, M.W., Yang, Z., Innes, L.M., Kandasamy, B., David, L.L., Yoshioka, C., and Shyng, S.L. (2019). Mechanism of pharmacochaperoning in a mammalian KATP channel revealed by cryo-EM. Elife 8.

Martin, G.M., Yoshioka, C., Rex, E.A., Fay, J.F., Xie, Q., Whorton, M.R., Chen, J.Z., and Shyng, S.L. (2017b). Cryo-EM structure of the ATP-sensitive potassium channel illuminates mechanisms of assembly and gating. Elife 6.

Nichols, C.G. (2006). KATP channels as molecular sensors of cellular metabolism. Nature 440, 470–476.

Olson, T.M., Alekseev, A.E., Moreau, C., Liu, X.K., Zingman, L.V., Miki, T., Seino, S., Asirvatham, S.J., Jahangir, A., and Terzic, A. (2007). KATP channel mutation confers risk for vein of Marshall adrenergic atrial fibrillation. Nat. Clin. Pract. Cardiovasc. Med. 4, 110–116.

Pipatpolkai, T., Usher, S., Stansfeld, P.J., and Ashcroft, F.M. (2020). New insights into KATP channel gene mutations and neonatal diabetes mellitus. Nat Rev Endocrinol 16, 378–393.

Proks, P., de Wet, H., and Ashcroft, F.M. (2010). Activation of the K(ATP) channel by Mg-nucleotide interaction with SUR1. J. Gen. Physiol. 136, 389–405.

Reimann, F., Tucker, S.J., Proks, P., and Ashcroft, F.M. (1999). Involvement of the n-terminus of Kir6.2 in coupling to the sulphonylurea receptor. J Physiol 518 (Pt 2), 325–336.

Smeland, M.F., McClenaghan, C., Roessler, H.I., Savelberg, S., Hansen, G.A.M., Hjellnes, H., Arntzen, K.A., Muller, K.I., Dybesland, A.R., Harter, T., et al. (2019). ABCC9-related Intellectual disability Myopathy Syndrome is a KATP channelopathy with loss-of-function mutations in ABCC9. Nat Commun 10, 4457.

Sung, M.W., Yang, Z., Driggers, C.M., Patton, B.L., Mostofian, B., Russo, J.D., Zuckerman, D.M., and Shyng, S.L. (2021). Vascular KATP channel structural dynamics reveal regulatory mechanism by Mg-nucleotides. Proc. Natl. Acad. Sci. U. S. A. 118.

Tucker, S.J., Gribble, F.M., Zhao, C., Trapp, S., and Ashcroft, F.M. (1997). Truncation of Kir6.2 produces ATP-sensitive K+ channels in the absence of the sulphonylurea receptor. Nature 387, 179–183.

van Bon, B.W., Gilissen, C., Grange, D.K., Hennekam, R.C., Kayserili, H., Engels, H., Reutter, H., Ostergaard, J.R., Morava, E., Tsiakas, K., et al. (2012). Cantu syndrome is caused by mutations in ABCC9. Am. J. Hum. Genet. 90, 1094–1101.

Wang, M., Wu, J.X., and Chen, L. (2022a). Structural Insights Into the High Selectivity of the Anti-Diabetic Drug Mitiglinide. Front Pharmacol 13, 929684.

Wang, M., Wu, J.X., Ding, D., and Chen, L. (2022b). Structural insights into the mechanism of pancreatic KATP channel regulation by nucleotides. Nat Commun 13, 2770.

Wu, J.X., Ding, D., and Chen, L. (2022). The Emerging Structural Pharmacology of ATP-Sensitive Potassium Channels. Mol. Pharmacol. 102, 234–239.

Wu, J.X., Ding, D., Wang, M., Kang, Y., Zeng, X., and Chen, L. (2018). Ligand binding and conformational changes of SUR1 subunit in pancreatic ATP-sensitive potassium channels. Protein Cell 9, 553-567.

Zhao, C., and MacKinnon, R. (2021). Molecular structure of an open human KATP channel. Proc. Natl. Acad. Sci. U. S. A. 118.

